# IRIS PIGMENTATION IRREGULARITIES FOLLOWING AN AVIAN INFLUENZA OUTBREAK: IMPLICATIONS FOR DISEASE SURVEILLANCE AND POPULATION MONITORING IN A COLONIAL SEABIRD

**DOI:** 10.1101/2025.10.31.685885

**Authors:** C. Petalas, J.A. Giacinti, J.F. Provencher, J.-F. Rail, S. Avery-Gomm, R.A. Lavoie, K.H. Elliott, Y. Seyer

## Abstract

Emerging infectious diseases can have catastrophic impacts on wildlife populations, yet identifying individuals that survived exposure, especially when external symptoms are absent, remains challenging. Since 2021, a virulent strain of highly pathogenic avian influenza virus (HPAIV H5N1 clade 2.3.4.4b) has caused unprecedented mortality in wild birds across continents. Northern Gannets (*Morus bassanus*) are among the species that suffered significant population declines in Europe and North America. At North America’s largest gannet colony (Bonaventure Island) dramatic mortality and reproductive failure occurred in 2022. Following this event, researchers noted a subset of gannets displaying irregular iris pigmentation, raising the possibility that this visible change may indicate a lasting effect of infection. Here, we build on earlier observations linking irregular iris pigmentation to HPAIV exposure in gannets using anti-nucleoprotein (NP) and anti-hemagglutinin (H5) antibodies. This provides the first quantitative test of this relationship using serological data and field-based digital photography. Iris irregularities were strongly associated (ρ = –0.72) with antibodies to NP, supporting the hypothesis that they can indicate past exposure. The likelihood of NP antibody detection increased with iris pigment irregularity—about 50% likelihood at 40% irregularity, 65% at 50%, 77% at 60%, and over 90% above 77% irregularity. Moderate correlations (ρ = 0.30) were observed for H5 antibodies. Our findings provide quantitative support for the hypothesis that iris pigmentation irregularities may serve as a visible, non-invasive marker of past HPAIV exposure in gannets. If validated across colonies and years, iris assessment could offer a rapid tool for tracking population health and recovery following HPAIV outbreaks, enhancing conservation monitoring and disease surveillance.

**Lay Summary:** Our research shows clear evidence that irregular iris pigmentation in Northern Gannets is strongly linked to past infection with a deadly bird flu strain. This visible marker could allow scientists to rapidly identify surviving birds and monitor population health after outbreaks, helping conservation efforts across affected colonies.

**Highlights:**

- Iris pigmentation irregularities reveal past avian influenza exposure.
- Extent of iris pigmentation irregularity correlates with antibody levels.
- Exposure likelihood increases with degree of iris pigmentation irregularity.
- Could provide a rapid, non-invasive tool for wildlife disease and conservation monitoring.

**Graphical abstracts:** 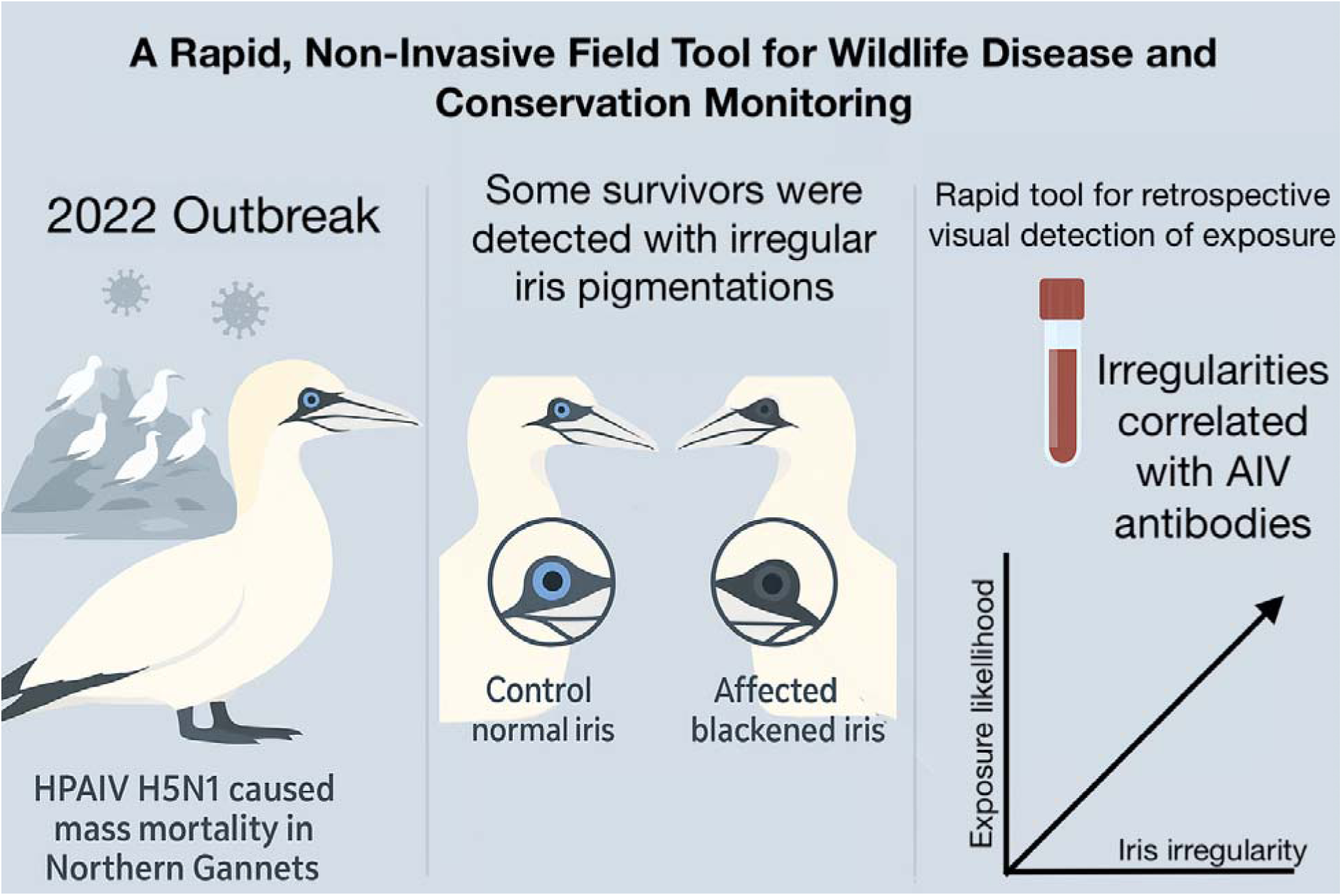

## 1. Introduction

Avian influenza viruses (AIVs) have circulated in some species of wild birds for over a century (Hoye et al. 2010; Lycett et al. 2019), but recent decades have seen an increase in both the frequency and severity of detected outbreaks (Stallknecht and Brown, 2008; Chatziprodromidou et al., 2018; Blagodatski et al., 2021). Since 2021, a particularly virulent strain, highly pathogenic avian influenza virus (HPAIV) H5N1, clade 2.3.4.4b, has caused unprecedented global mortality in wild birds and poultry (Camphuysen et al., 2022; Wille and Barr, 2022). Seabirds have been heavily affected, likely due to factors such as limited prior immunity, dense nesting conditions, and high rates of direct contact, all of which facilitate viral transmission (Lang et al., 2016; Dewar et al., 2023; McPhail et al., 2024). Mass die-offs due to HPAIV have been documented at several seabird breeding colonies, with some populations experiencing losses exceeding 70% (Banyard et al., 2022; Camphuysen et al., 2022; Rijks et al., 2022; Pohlmann et al., 2023; Knief et al., 2024; Tremlett et al., 2024). HPAIV has therefore emerged as a major conservation concern for some seabird species, contributing to one of the largest documented wild bird mortality events associated with avian influenza (CMS-FAO, 2022).

Northern Gannets (*Morus bassanus*) have been among the most severely impacted seabird species, with large-scale mortality events reported at 40 of 53 global colonies in 2022, including five of six Canadian colonies (Avery-Gomm et al., 2024; Lane et al., 2023, Seyer and Guillemette, 2023). Their dense colonial nesting conditions likely facilitated rapid viral spread (McPhail et al., 2024), as well as limited prior immunity to AIV (McLaughlin et al., 2025). Since 2022, HPAIV has caused mass mortality events at most gannet colonies across the North Atlantic, raising urgent questions about colony-level vulnerability, immunity, and the potential for repeated outbreaks. Long-term monitoring has documented cumulative environmental stressors for gannets, including prey depletion, ocean warming, and anthropogenic activities (Guillemette et al., 2018; Grémillet et al., 2020; Montevecchi et al., 2021). In 2022, gannets experienced high HPAIV-related mortality (Jessopp et al., 2023; Giacinti et al., 2024a), adding to existing conservation concerns and highlighting the importance of understanding not only short-term survival but also long-term recovery potential in gannet populations worldwide.

While direct mortality has dominated attention, emerging evidence suggests HPAIV may also have sublethal effects on survivors, including altered movement patterns, reduced reproductive success, and impaired condition (Duriez et al., 2023; Careen et al., 2024; Lewis et al., 2025). Detecting these impacts, however, remains difficult, particularly in remote colonies where invasive sampling is challenging. That difficulty has prompted interest in developing non-invasive field tools that can help identify individuals previously exposed to HPAIV and support monitoring of colony recovery and resilience following mass mortality events (Nalepa et al., 2024).

One promising field indicator is the recent observations of iris pigmentation irregularities, typically partial or complete darkening of the otherwise pale-blue iris, in gannets that survived the outbreak (Lane et al., 2023). While gannet irises are naturally pale-blue, changes in iris pigmentation may result from melanin deposition or inflammatory processes that alter the appearance of the iris (Corbett et al., 2024). Alteration of pigments, whether via immune-mediated mechanisms or tissue inflammation during infection, can lead to visible darkening of the iris (Toomey et al., 2010; Olson and Owens, 2005). Although the physiological basis and functional consequences of this iris darkening remain unknown, these mechanisms suggest a plausible link between HPAIV exposure and iris appearance, although it remains uncertain whether this response is specific to HPAIV or reflects a broader inflammatory reaction. Ocular manifestations have also been reported in other large birds that tested positive for H5 antibodies following an HPAIV outbreak (Alexandrou et al., 2025), suggesting that AIV infection can affect the eyes of multiple species, though the specific symptoms may vary.

If reliably associated with AIV antibody presence, iris pigmentation irregularities could serve as a rapid, practical screening tool to identify previously exposed individuals and to estimate rates of sublethal infection in the population. Applied at the colony level, repeated assessments of iris pigmentation could track the prevalence of prior infection over time, providing a non-invasive means to gauge post-outbreak survival and recovery. Linking these data with reproductive metrics would allow evaluation of how disease exposure influences population resilience, thereby integrating health surveillance into conservation monitoring.

Lewis et al. (2025) recently demonstrated the potential of iris pigmentation irregularities as a non-invasive tool for post-outbreak population monitoring, showing that gannets with black irises had similar breeding success to unaffected individuals. However, the physiological link between iris pigmentation and immune status has not been empirically validated. No studies have yet tested whether iris pigmentation irregularities correspond with serological markers of HPAIV exposure in gannets, a critical gap for confirming the utility of this trait as a monitoring tool.

Addressing this question is essential for developing effective, non-invasive approaches to track the longer-term impacts of HPAIV on seabird populations and inform conservation strategies. Here, we directly test this link by combining serological and photographic data to assess whether iris pigmentation irregularities are associated with antibody-based evidence of past AIV exposure. Using whole blood collected on filter strips, an efficient, field-adaptable method for serological sampling (Giacinti et al., 2025), we tested for anti-nucleoprotein (NP) and anti-hemagglutinin (H5) antibodies which provide evidence of past exposure to AIV in wild birds (Giacinti et al., 2025). We also developed a standardized method to quantify iris pigmentation irregularities from digital photographs and compared antibody levels between gannets with and without visible irregularities during the 2023 breeding season.

We hypothesize that variations in iris pigmentation irregularities correspond to differences in antibody response, reflecting individual variation in exposure intensity or immune activation. Specifically, we predicted that individuals with more irregular (darker) irises would exhibit stronger antibody signals and would be more likely to test positive for anti-NP and anti-H5 based on established thresholds. Additionally, we evaluated whether antibody presence was associated with body mass, a proxy for physiological condition, based on previous findings that viral infection and immune activation can reduce condition in seabirds (Siebert et al., 2012) and may impair vision and, consequently, foraging efficiency. Because individuals exhibit varying degrees of iris irregularity, we further hypothesized that pigmentation irregularities may decrease over the breeding season, reflecting possible fading of the extent of irregularity with time. This study aims to evaluate whether iris pigmentation could serve as a practical, low-cost field marker of prior HPAIV exposure, thereby supporting efforts to monitor avian influenza impacts in wild seabird populations.

## 2. Materials and Methods

### 2.1. Sample collection

Canada supports the entire North American breeding population and ∼13% of the global gannet population (Mowbray, 2020), including the largest colony (∼104,000 breeding individuals, Rail, 2021) located on Bonaventure Island, Québec, Canada (48.4943° N, 64.1608° W) in the Gulf of St. Lawrence. Bonaventure Island is located in the Parc national de l’Île-Bonaventure-et-du-Rocher Percé and is protected under provincial (a park under the Société des établissements de plein air du Québec; Sépaq) and federal (a Migratory Bird Sanctuary under Environment and Climate Change Canada; ECCC) jurisdictions. Gannets breed on the south-east side of the island on cliffs or on the plateau where the majority of pairs nest (Figure 1). On the plateau, nests are regularly spaced ∼60-80 cm apart (Poulin, 1968). From 11 to 14 September 2023, during the breeding season, 65 adult gannets were captured on or near their nests at the Bonaventure Island colony.

**Figure 1.**
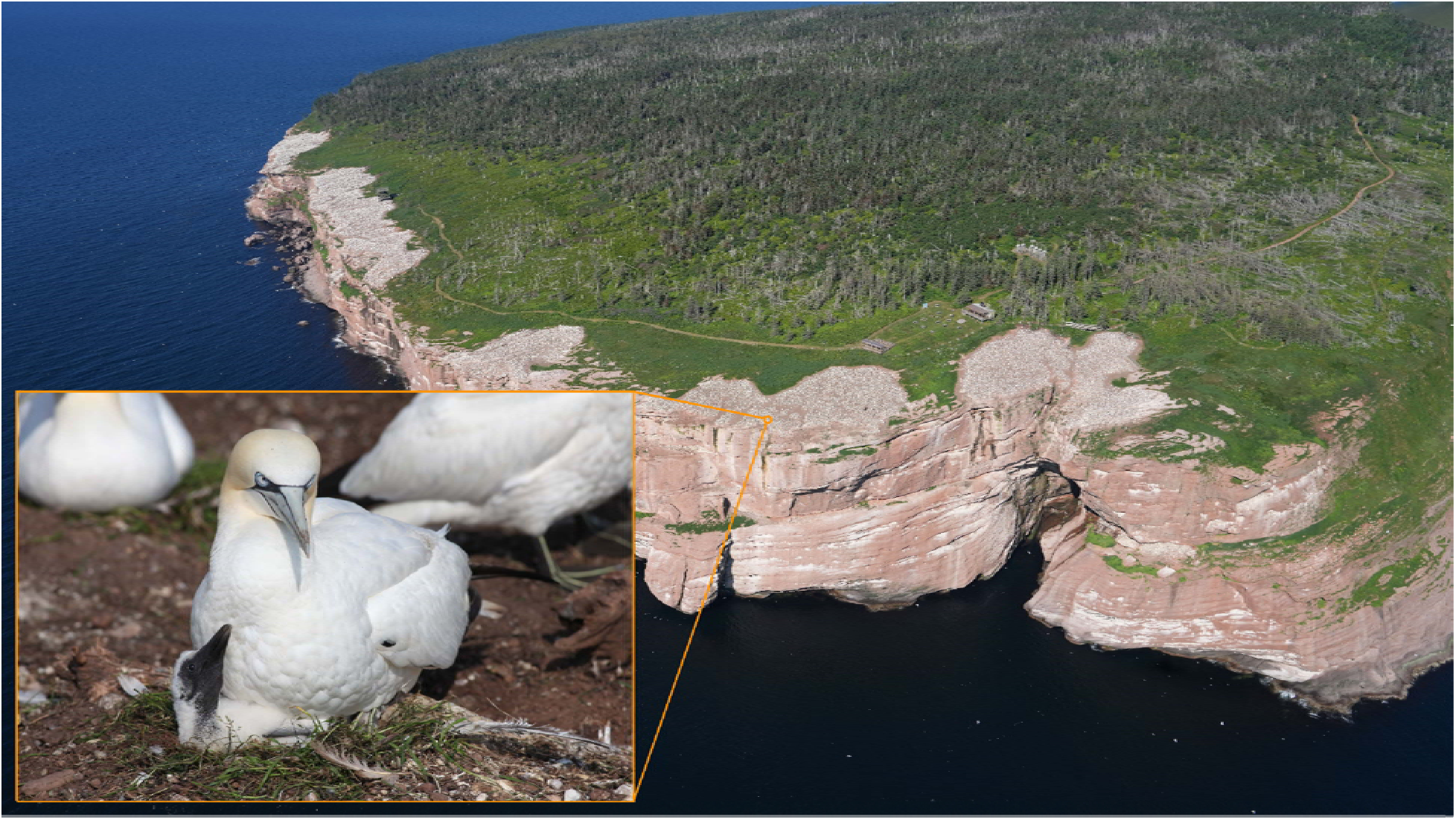
Northern Gannet (*Morus bassanus*) breeding colony at Bonaventure Island, Québec. The inset shows an adult gannet with its chick at the nest. Photographs by J.F.R.

Breeding gannets were captured across the plateau area at the outer edge of the colony while on or near their nest using a noose-pole. Individuals were targeted based on iris pigmentation, attempting to sample both birds with irregular and regular iris pigmentation. Of the 30 individuals targeted for having irregular iris pigmentation, 13% were captured on nests with chicks; the remaining birds were either on empty nests (47%) or not on a nest at the time of capture (40%). In contrast, 80% of the 35 individuals with regular iris pigmentation were captured on active nests with chicks, 17% on an empty nest and 3% not on a nest. All individuals appeared in good body condition and exhibited no overt behavioral impairments (e.g., seemed to be regularly provisioning or conducting foraging activity). In addition to the methods described below related to this study, all captured birds were marked with a United States Geological Survey (USGS) metal band. All live bird sampling followed approved animal use protocols, applicable federal and provincial wildlife permits, and safe work procedures.

The tarsal vein was pricked using a 24-gauge needle. Approximately 3–4 drops of whole blood (∼0.1 mL) were collected from each bird directly from the vein onto a Nobuto blood filter strip (Advantec MFS, Inc., USA; Product #49010010), fully saturating the strip following the procedure described in Giacinti et al., (2025). After sampling, all filter strips were air-dried in tented paper coin envelopes for at least 24 hours and preserved at room temperature.

Oropharyngeal and cloacal swabs were also taken from each bird to test for HPAIV infection. The oral and cloacal swabs were placed into a single vial of 3 mL Universal Transport Medium (Copan Italia S.p.A., Brescia, Italy) and represent a single sample per individual. Each tube was immediately placed in a freezer bag for temporary storage during daily field activities and stored at −20°C within a few hours until they could be transported to the laboratory for avian influenza testing.

During the handling of adult gannets, photographs of each individual’s left and right eyes were captured using a handheld camera approximately 20 cm away from the individual without using the flash. Date and time information was recorded for each photograph. Gannets were held by the bill and body in a consistent manner. Researchers attempted to capture photographs under similar bright light conditions (collections conducted over four days at the same times of day) to ensure consistent illumination for all photographs and pupil diameter measurements. Prior to release, the body mass of each bird was measured using a 4000-g Pesola scale.

Moreover, during the 2024 breeding season (July to September), we selected 7 individual gannets with iris pigment irregularities for repeated iris pigmentation assessments to assess within-individual variability over time. Each bird was photographed multiple times from afar throughout incubation and chick-rearing stages.

### 2.2. Sample preparation and laboratory testing

The detection of anti-NP antibodies was performed using the IDEXX AI MultiS Screen Ab test (IDEXX Canada, Product # 99-12119) at the National Wildlife Research Center (NWRC). Nobuto filter strips saturated with whole blood were cut and eluted in 400 μl of phosphate-buffered saline (PBS; pH 7.2; GibcoTM, USA, Product # 20012027) at 4°C for 24 hours (as described by Giacinti et al., 2025). ELISA testing was then conducted according to the manufacturer’s instructions (Brown et al., 2009). All eluates were also tested for anti-H5 antibodies at the National Centre of Foreign Animal Disease (NCFAD), using previously established methods (Hochman et al., 2023). The sample-to-negative (S/N) ratio and percent inhibition (PI%) values derived from these semi-quantitative ELISA assays provide relative measures of antibody binding to influenza antigens, where lower S/N ratios and higher PI% values indicate stronger binding to the assay antigen.

An optimized threshold of S/N < 0.77 was used to classify samples as positive for anti-NP antibodies, following threshold optimization performed at NWRC for Nobuto strip testing as discussed in Giacinti et al., (2025). A PI value ≥ 20.37% was considered positive for H5 antibodies, based on the optimized cutoff for Nobuto strips (Giacinti et al., 2025). These assays detect antibodies broadly reactive to influenza A, but do not distinguish HPAIV from low-pathogenic H5 strains.

Pooled oropharyngeal and cloacal swabs were submitted to the Animal Health Laboratory (AHL) at the University of Guelph, where total RNA was extracted using the QIAamp Viral RNA Mini Kit (Qiagen). Samples were tested using real-time reverse-transcription PCR (RT-PCR) targeting the influenza A matrix (M) gene (Nagy et al., 2021) and a specific H5 assay (James et al., 2022), following standardized Canadian Animal Health Surveillance Network protocols (Giacinti et al., 2024a). Any samples testing positive at AHL would have been forwarded to the NCFAD for confirmatory testing, including additional RT-PCR, virus isolation, and sequencing. However, all samples in this study tested negative.

### 2.3. Image processing

All eye images were processed manually by the same individual to ensure reproducibility and consistency using Adobe Photoshop version 25.4. Each image was first cropped to isolate the iris, excluding the pupil and sclera. If necessary, to standardize the visual data, brightness was adjusted using the “Brightness/Contrast” tool to improve clarity. The images were then converted to grayscale using the “Mode” function, and subsequently transformed into a binary (black-and-white) format using the “Threshold” adjustment tool (Figure 2). This resulted in only black and white pixels while removing all color. The degree of irregular iris pigmentation was quantified by calculating the ratio of black to white pixels within the isolated iris area. This ratio, hereafter is referred to as iris pigmentation irregularity (%), represents the extent of visible dark pigmentation within the iris, quantified on a scale from 0% (no irregularity) to 100% (complete darkening). Iris pigmentation irregularities were evaluated for each eye separately, and for analyses, the eye exhibiting the most pronounced pigmentation irregularity (hereafter, iris with greater irregularity) was used to assign an individual iris score.

**Figure 2.**
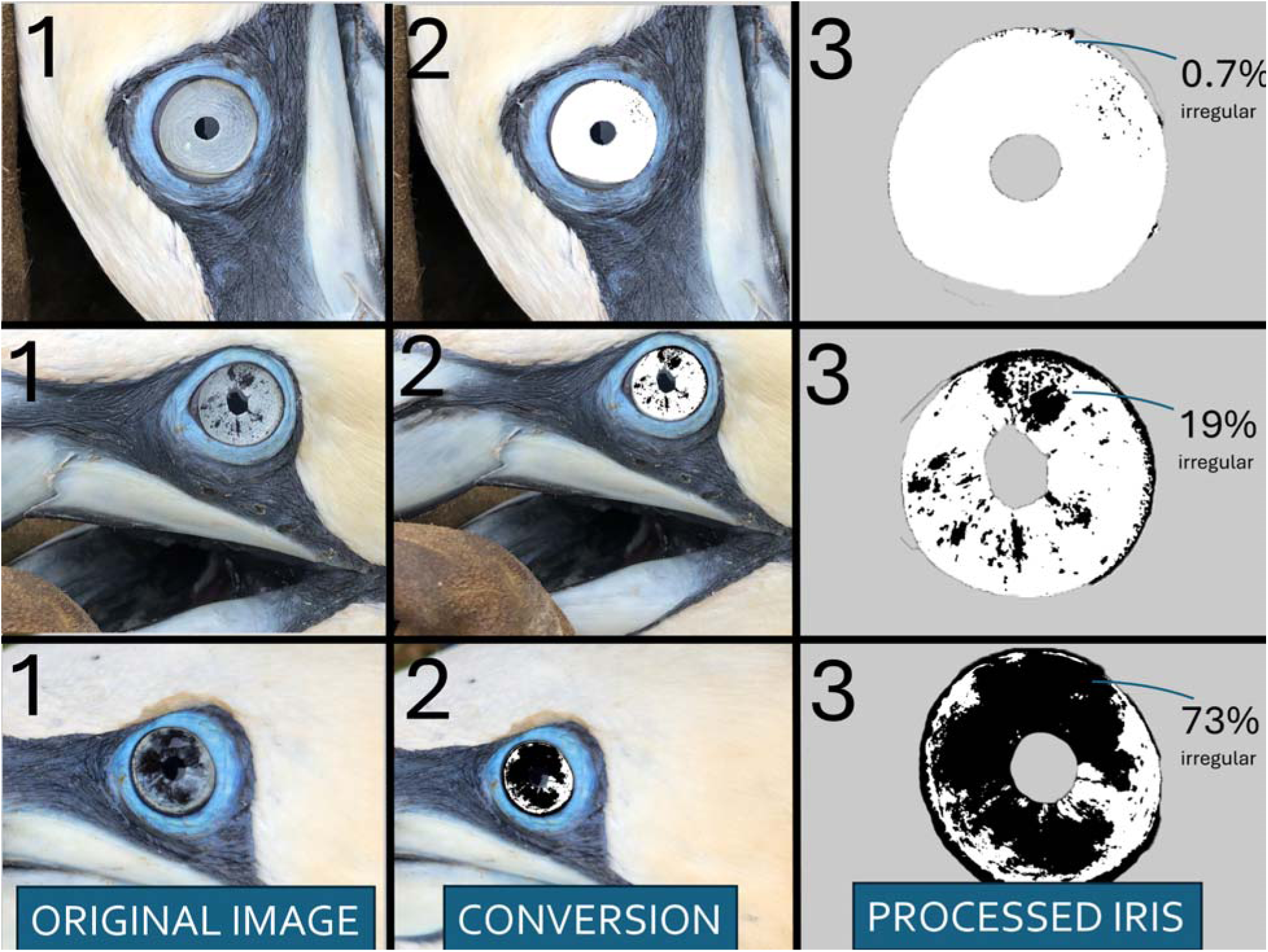
Example of image processing procedure where Northern Gannet photographs taken in the field (step 1) were analysed using Photoshop (step 2) to isolate the iris and then calculate the percentage of irregularities in each iris (step 3).

### 2.4. Quality assurance and quality controls (QA/QC)

To ensure the consistency of image processing, we implemented a quality assurance protocol. For 10% of the individuals, we randomly duplicated the image processing and compared the results for consistency. Additionally, for 5% of the individuals, we triplicated the image processing. Because traditional relative percent difference (RPD) calculations can produce exaggerated values when measurements are near zero, we used a modified RPD to reduce this sensitivity:

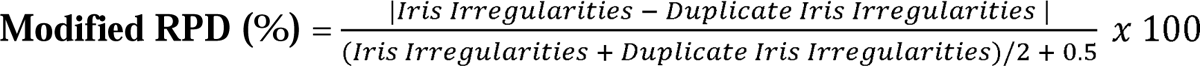

A constant of 0.5 was added to the denominator based on the distribution of our data. An RPD < 20% was considered acceptable. A modified RPD was used to reduce inflation of error at low values, resulting in an average RPD of 16.20 ± 17.42%, with 73.10% of values falling within the pre-established acceptable range. In addition, we calculated the absolute difference between duplicate measurements. Small absolute differences, despite potentially high relative percent differences (RPD), were considered acceptable when they fell within a pre-established threshold (<0.5%), with an average error of 0.55 ± 3.86%, supporting consistency even when RPD values were high due to near-zero measurements. A linear regression model was fit to assess how much variation in the duplicate measurements could be explained by the original measurements. A strong correlation between original and duplicate values (R² = 0.993) further confirmed measurement precision, meaning that the original measurement explained 99.30% of the variance in the duplicated measurement. This strong model fit suggests that repeated processing of the same images yields nearly identical results, suggesting that the original measurements are highly repeatable and not substantially influenced by random error or subjectivity in image processing.We can be confident that the original measurements accurately reflect true variation in iris pigmentation irregularity among individuals rather than artifacts of the measurement procedure.

To establish a baseline threshold for iris pigmentation irregularity in gannets prior to the outbreak, we processed images of gannets taken from the same Bonaventure Island colony before 2021. These archive images were taken from random individuals over multiple years without following a standardized procedure. They were taken with a Canon EOS 5D Mark II or Canon EOS 10D and a Canon 75-300 or 400 mm lens from a distance. These images, spanning from 2004 to 2018, provided a reference baseline for the typical range of iris irregularities observed in the population before the outbreak (n = 28). To evaluate whether birds sampled post-outbreak but classified as having “regular” irises showed comparable pigmentation, we compared this baseline to individuals sampled in 2023.

### 2.5. Statistical analysis

All analyses were conducted using R software version 4.4.2 (R Core Team, 2025). To assess the relationship between iris pigmentation irregularities and antibody levels, we conducted two sets of analyses. First, we used Spearman’s rank correlation tests using the *cor.test()* function in R to examine associations between iris pigment irregularities and the continuous ELISA test results: S/N ratios for anti-NP and PI% for anti-H5. These were tested separately for anti-NP and anti-H5. These non-parametric correlations were appropriate due to the continuous nature of the antibody measures and the non-normal distribution of iris pigment irregularity scores. For each bird, we calculated correlations using two iris metrics: (1) the average irregularity across the left and right irises and (2) the irregularity score of the iris with greater irregularity. Results are reported as Spearman correlation coefficients (ρ) with corresponding p-values for each antibody type (anti-NP and anti-H5). We also used the same approach to test whether the body mass of the individual was correlated with the continuous antibody results (anti-NP and anti-H5).

Secondly, to assess whether iris pigment irregularity was predictive of AIV serological status, we conducted logistic regressions using the *glm()* function in R with a binomial error distribution. The outcome variable was the binary interpretation of the anti-NP antibody test (“Positive” or “Negative”). Iris pigment irregularity was included as the predictor in two separate models: one using the average across both irises and another using the iris with greater irregularity. Parallel logistic regressions were conducted using binary anti-H5 antibody status (based on established PI% thresholds) as the outcome.

To assess whether anti-NP and anti-H5 antibody levels differed among birds with varying degrees of iris pigment irregularity, we categorized individuals based on their average iris pigment irregularity score (across both irises) into three groups: Low (<33%), Moderate (33– 66%), and High (>66%). Given that assumptions of normality and homogeneity of variance were not met for the antibody data, we used non-parametric Kruskal-Wallis tests to compare antibody levels across these groups. Post-hoc pairwise comparisons were conducted using Wilcoxon rank-sum tests with Bonferroni correction to identify specific group differences.

To explore whether iris pigmentation irregularities may affect physiological condition via impaired vision and reduced foraging efficiency, we conducted an analysis examining body mass across iris pigment irregularity scores (low, moderate, high). Assumptions of normality and homogeneity of variances were assessed using the Shapiro-Wilk test and Levene’s test, respectively. Due to violation of homogeneity of variances, a non-parametric Kruskal-Wallis rank sum test was used.

All figures were created using *ggplot2*. We used a significance threshold of 0.05. Means are presented with standard deviation (± SD).

## 3. Results

No mass mortality events were reported at Bonaventure Island during the 2023 breeding season, and all birds sampled in 2023 tested negative for active HPAIV infection via RT-PCR targeting the matrix gene and H5 subtype (n = 65). Of these, 40% tested positive for anti-NP antibodies. Among NP-positive birds, 85% exhibited high or moderate iris pigmentation irregularities (Table 1). 20% tested positive for anti-H5 antibodies. Among these, 54% had high iris pigmentation irregularities and 46% had low pigmentation; none fell into the moderate category (Table 1).

**Table 1.**
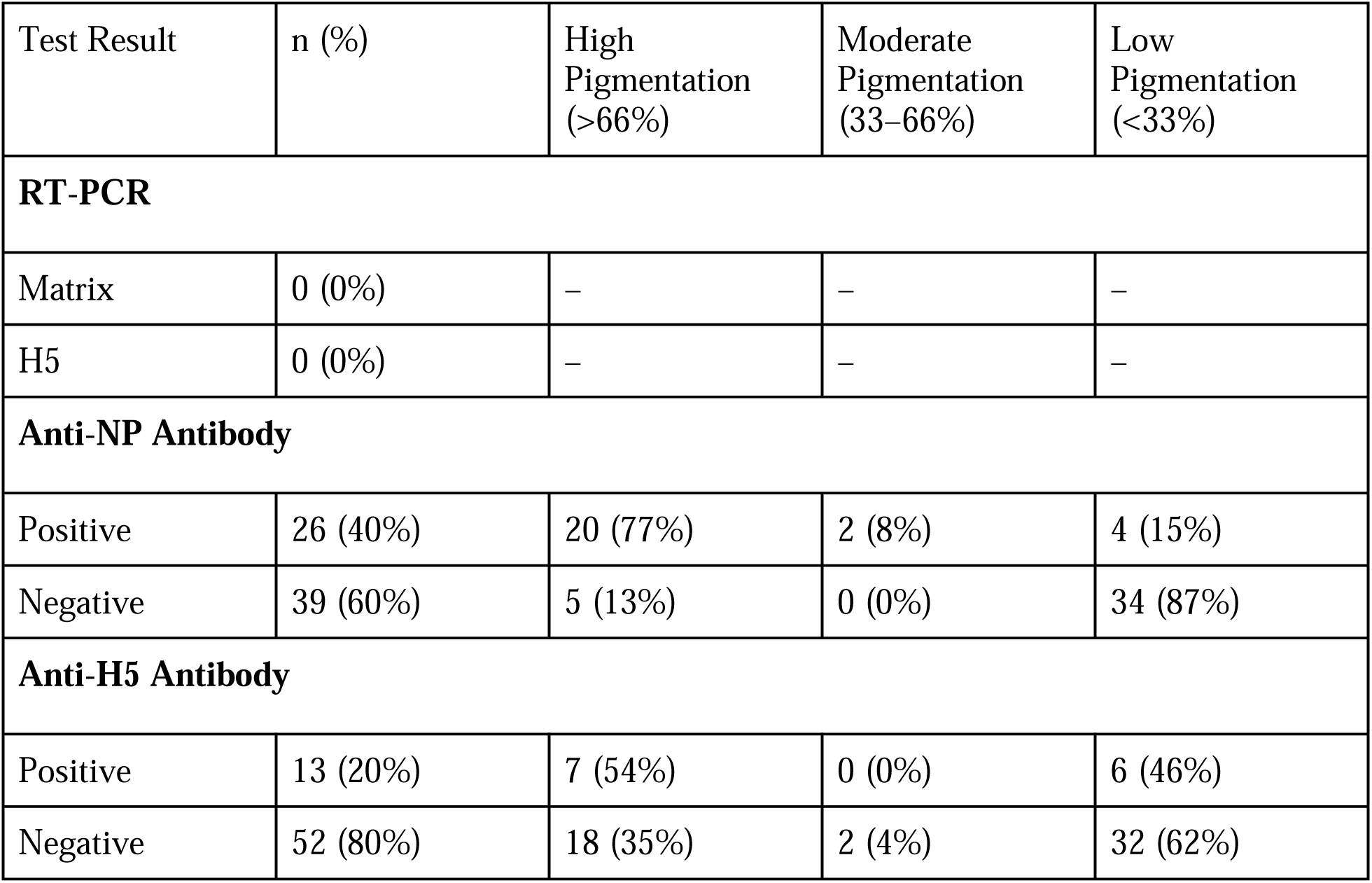
Summary of HPAIV diagnostic results and iris pigmentation categories in 65 Northern Gannets sampled at Bonaventure Island in 2023.

The pre-outbreak baseline for iris pigmentation irregularities was 4.39 ± 1.84%, with a range of 0.22–8.02%. Birds sampled post-outbreak in 2023 and classified as having “regular” irises had a mean of 2.52 ± 2.40%, with a range of 0.09–11.76%. This comparison confirmed that birds without visible pigmentation changes post-outbreak generally fell within the normal pre-outbreak range.

Results are shown for average iris irregularity only, as model coefficients and significance were consistent when using the iris with greater irregularity (Supplemental Figures S1-S3). Iris pigment irregularity was negatively associated with anti-NP S/N values (ρ = –0.72, *p* < 0.001; Figure 3). This suggests that birds with greater iris pigmentation irregularity tended to have lower S/N values - i.e., more positive results for anti-NP antibodies.

**Figure 3.**
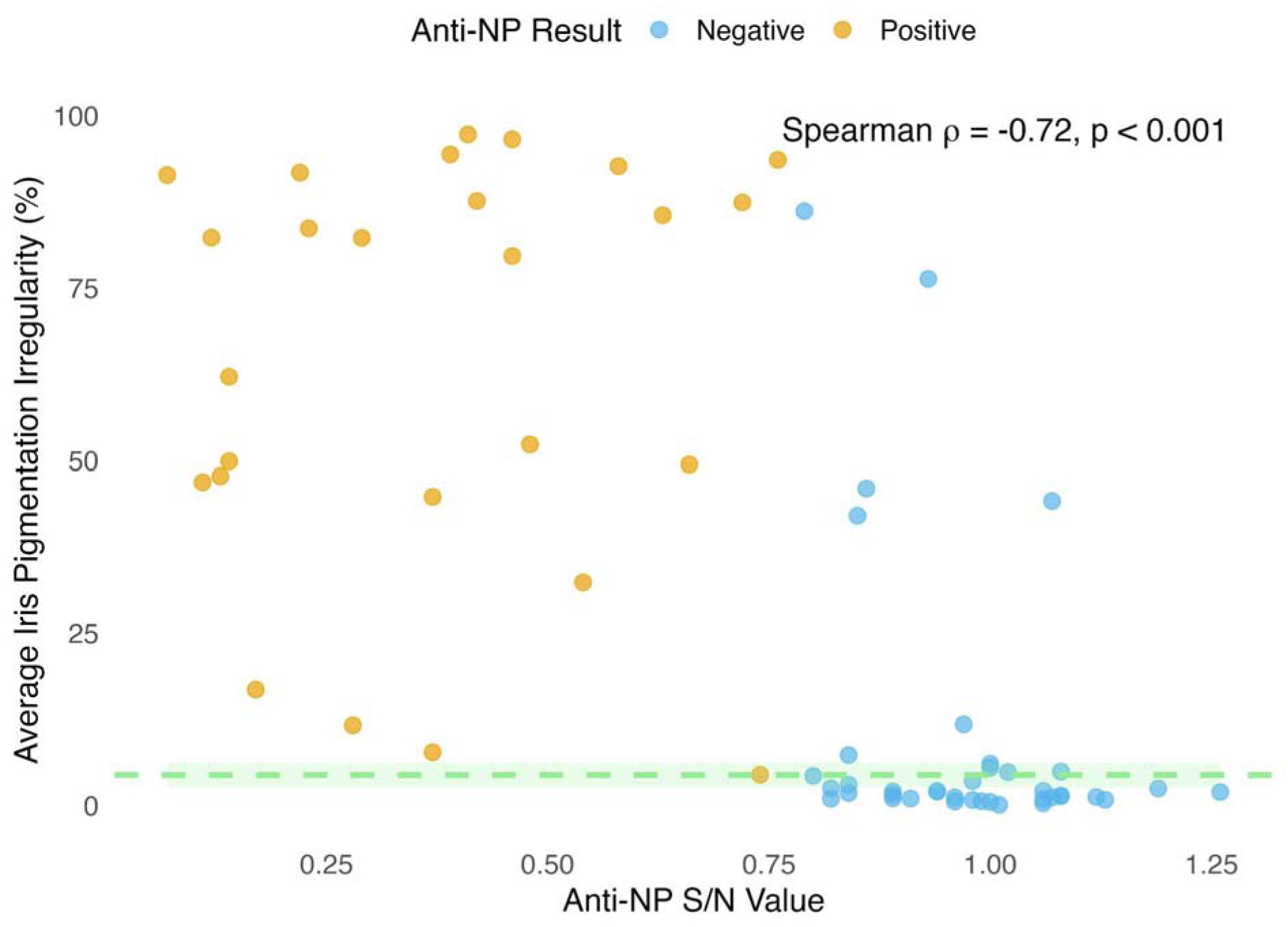
Correlation between anti-NP S/N values and iris pigmentation irregularity (%) for average irregularity across both irises. Points are colored by anti-NP result interpretations (“Positive” or “Negative”). The dashed green line shows the average pre-outbreak (pre-2021) irregularity, with the shaded band representing ±1 standard deviation. Note that lower S/N ratios correspond to stronger anti-NP antibody detection. An optimized threshold of S/N < 0.77 was used to classify samples as positive for anti-NP antibodies, based on threshold optimization for Nobuto strip testing at NWRC (Giacinti et al., 2025).

There was a significant, though weak, positive correlation between average iris pigment irregularity and anti-H5 PI% (ρ = 0.28, *p* = 0.026; Figure 4), indicating that individuals with more irregular irises tended to have higher PI% - i.e., more positive results for anti-H5 antibodies.

**Figure 4.**
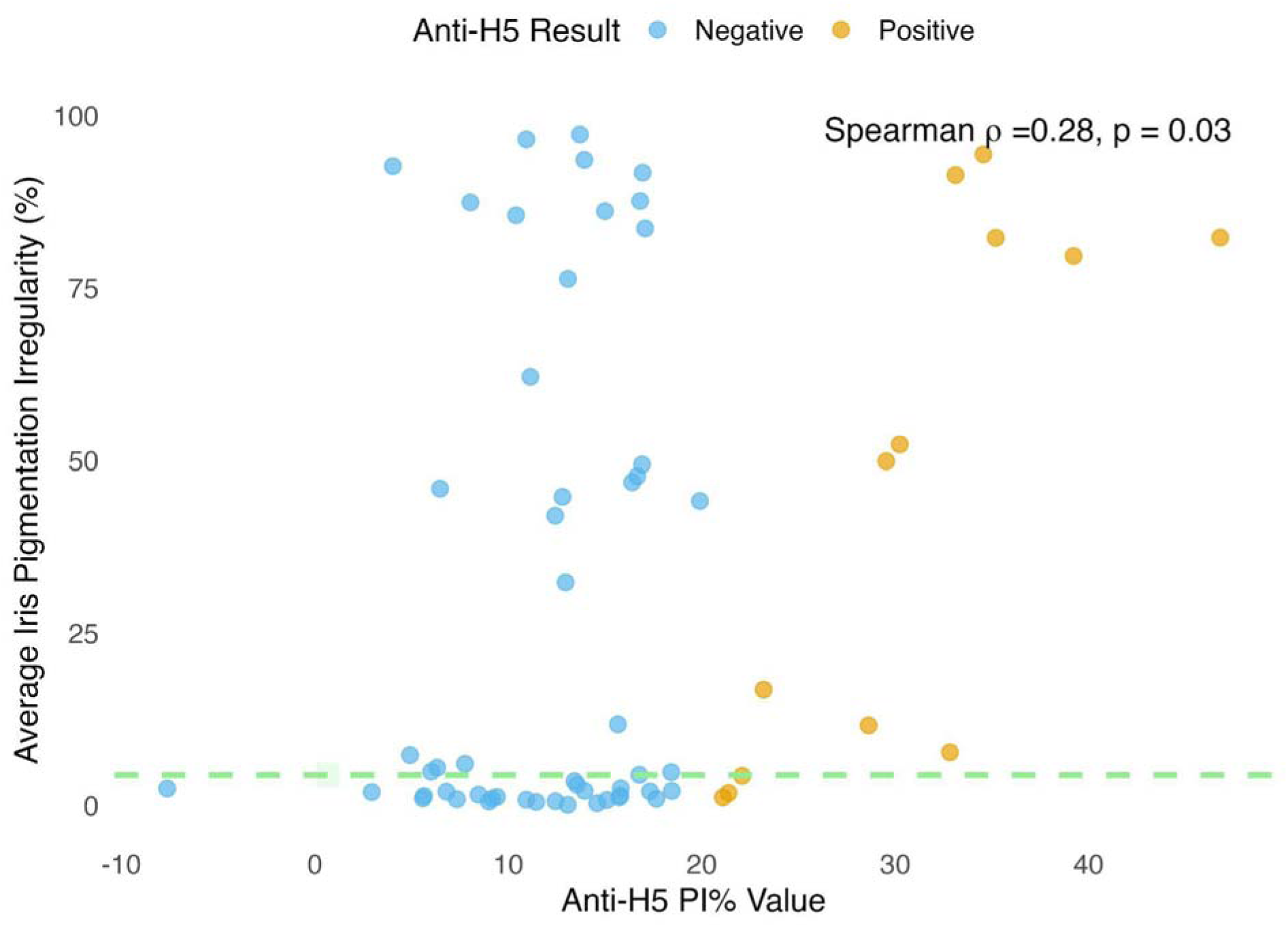
Correlation between anti-H5 PI% antibody detection and iris pigmentation irregularity (%) for average irregularity across both irises. Points are colored by anti-H5 result interpretations (“Positive” or “Negative”). The dashed light green line shows the average pre-outbreak (pre-2021) irregularity, with the shaded band representing ±1 standard deviation. Note that higher PI values correspond to stronger anti-H5 antibody detection. A PI value ≥ 20.37% was considered positive, reflecting the optimized cutoff for Nobuto strips (Giacinti et al., 2025).

There was a significant positive association between average iris pigment irregularity and the binomial probability of testing positive for anti-NP antibodies (β = 0.059, z = 4.58, *p* < 0.001; Figure 5). Specifically, for each 1% increase in average iris pigment irregularity, the odds of testing positive for anti-NP antibodies increased by approximately 6% of pigment irregularity.

**Figure 5.**
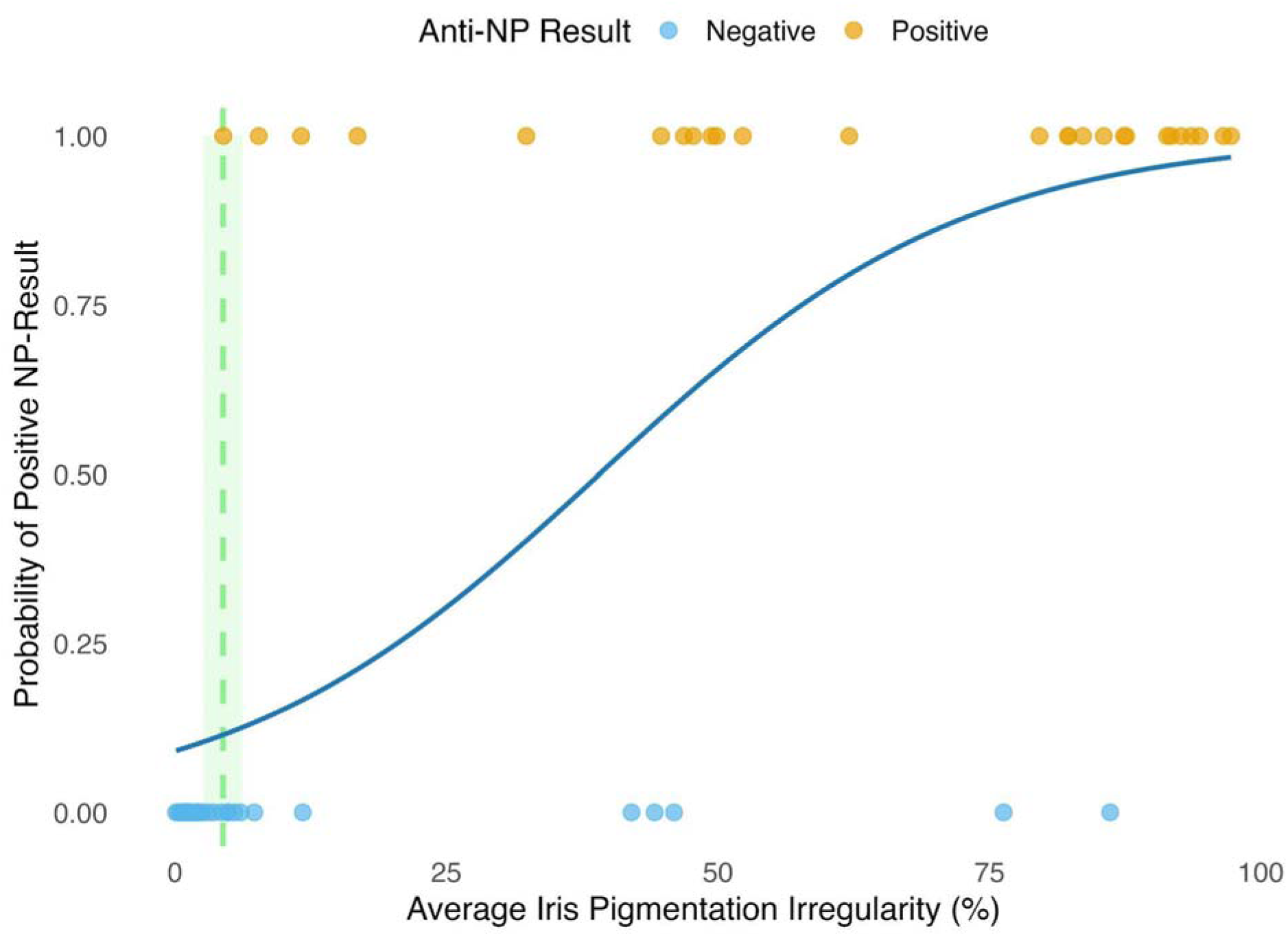
Logistic regression models predicting the probability of anti-NP S/N antibody positivity based on average iris pigment irregularity. Points are colored by test result interpretation (“Positive” or “Negative”) for each sample. The dashed light green line is average pre-outbreak (pre-2021) calculated irregularity, with the shaded band representing ±1 standard deviation. The solid blue line indicates the predicted probability of seropositivity based on the fitted regression model.

Average iris pigment irregularity was not a significant predictor of anti-H5 antibody status (β = 0.01, z = 1.39, *p* = 0.17). This suggests that variation in iris pigmentation was not associated with the probability of testing positive for anti-H5 antibodies.

There was a significant difference in anti-NP antibody values across birds with different levels of iris pigment irregularity based on average irregularity categorization (Kruskal-Wallis χ² = 27.15, df = 2, *p* < 0.001; Figure 6a). Birds with “high” iris pigment irregularity had significantly higher anti-NP antibody levels than those with “low” irregularity (*p* < 0.001), and birds with “moderate” irregularity also differed significantly from those with “low” irregularity (*p* = 0.003). However, no significant difference was observed between the “moderate” and “high” groups (*p* = 1.0).

**Figure 6.**
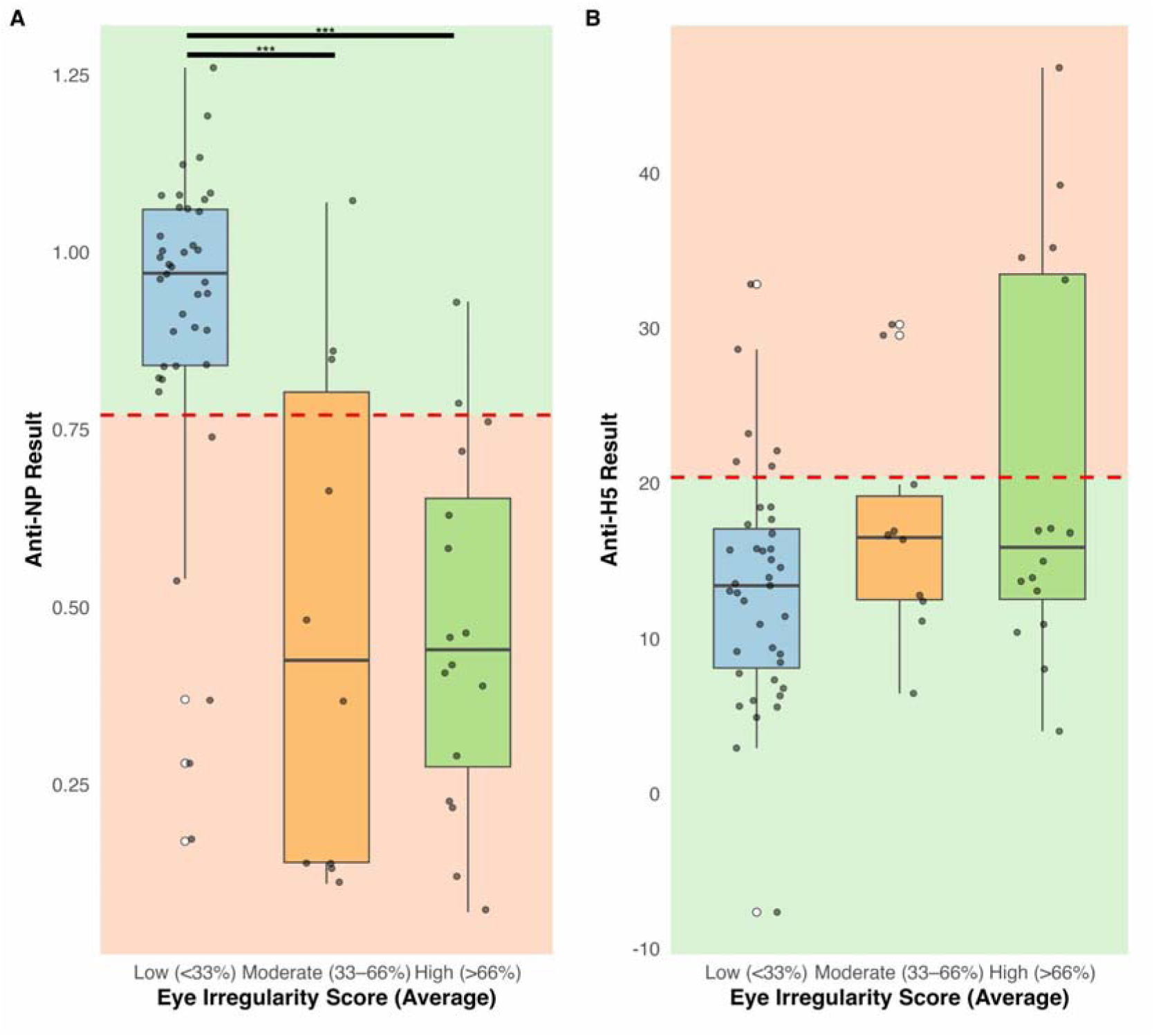
Boxplots of anti-NP antibody (A) and anti-H5 antibody (B) results across categories of iris pigment irregularity severity, classified based on the average irregularity score per bird (across both irises): Low (<33%), Moderate (33–66%), and High (>66%). In (A), the area above the threshold of 0.77 (colored in green) indicates negative results, and the area below the threshold (colored in red), indicates positive results and represents the optimized threshold value when using Nobuto filter strips on the IDEXX NP assay at NWRC (Giacinti et al. 2025). In (B), the area below the threshold of 20.4 (colored in green) indicates negative results, and the area above the threshold (colored in red), indicates positive results and represents the optimized threshold value when using Nobuto filter strips on the H5 assay at NWRC (Giacinti et al. 2025) *** indicates significant differences between groups (*p* < 0.0001).

Analysis of anti-H5 antibody values did not differ significantly among average iris pigment irregularity categories (χ² = 4.02, df = 2, *p* = 0.134; Figure 6b).

We did not find any correlation between body mass and anti-NP S/N values nor anti-H5 PI% (ρ = −0.13, *p* = 0.30; ρ = 0.23, *p* = 0.06, respectively). There were no significant differences in body mass among iris pigmentation categories (χ² = 2.81, df = 2, *p* = 0.25), suggesting that iris pigmentation irregularity level was not associated with variation in body condition.

Similar to 2023, there were no mass mortality events at the colony in 2024. Temporal assessment of 7 individual gannets revealed overall stable iris pigmentation across the 2024 breeding season (Figure S4). On average, the within-individual range in pigment irregularities was 4.6 ± 2.7%. Changes were not consistent in direction, 4 showed small decreases, while the remainder exhibited minor increases.

## 4. Discussion

### 4.1. Iris Pigmentation and Immune Response

This study provides an original methodological contribution to disease ecology by presenting evidence of a sublethal physiological change in gannets associated with AIV exposure. Iris pigmentation irregularities were first documented in gannets that survived the 2022 HPAIV outbreak (Lane et al., 2023). Previous work at Bonaventure Island also reported a shift in yolk anti-AIV antibody profiles following the outbreak (McLaughlin et al., 2025).

Building on those findings, we demonstrate that iris pigmentation irregularity in adult gannets is significantly associated with anti-NP antibody detection, suggesting that this visible trait may serve as a non-invasive marker of prior AIV exposure. This provides further evidence that some individuals survive AIV, including H5 variants, with important implications for assessing disease resilience and informing post-outbreak monitoring of population recovery.

Birds exhibiting increased departures from regular iris pigmentation (traces of black spots or almost fully blackened irises) were associated with the likelihood of detecting anti-AIV antibodies, including anti-H5. The presence of iris pigmentation irregularities in some antibody-negative birds, however, highlights the need for further validation of iris pigment changes as a reliable diagnostic tool.

The observed positive association between iris pigmentation irregularities and antibody reactivity, characterised by lower S/N values (for anti-NP) and higher PI% values (for anti-H5), was linked to more pronounced pigmentation irregularities. These findings suggest a potential biological link between immune response strength to AIV and ocular change, possibly driven by immune-mediated inflammation, viral damage to ocular tissue, or systemic immune processes affecting pigmentation. Though the underlying mechanisms remain unclear, evidence from other avian species (e.g., Yamamoto et al., (2016) demonstrated corneal opacity in ducks experimentally infected with H5N1 HPAIV, and Alexandrou et al., (2025) found that dalmatian pelicans (*Pelecanus crispus*) with cloudy corneas had H5-specific antibodies) highlights the potential capacity of AIV to induce ocular changes. Interestingly, other pathogens have been linked with changes in ocular structures; for example, *Toxoplasma gondii* has been shown to produce retinal lesions and ocular atrophy in canaries (Williams et al., 2001). The link between ocular irregularities and immune responses is an area that should be explored further. Post-mortem and histological analyses would help clarify these physiological pathways.

### 4.2. Stability and Temporal Dynamics of Pigmentation

Anti-NP antibody presence (a general marker of past AIV exposure) increased with iris pigmentation irregularity. Individuals with ∼40% irregularity had a >50% probability of testing positive for anti-NP antibodies, rising to ∼77% at 60%, ∼86% at 70%, and >90% at 80% (Figure 5). These conservative thresholds suggest that even moderate darkening should be considered a red flag for potential previous AIV exposure in gannets. Importantly, because control data from prior to the 2022 outbreak showed no comparable moderate or high iris pigmentation irregularity levels, it is reasonable to infer that pigmentation levels beyond control reflect physiological changes associated with the HPAIV outbreak. Nonetheless, long-term and comparative data are needed to validate these thresholds across years and populations and to assess whether similar ocular changes may occur in response to other pathogens, which would influence the specificity of this indicator.

We found evidence for the persistence of iris pigmentation changes, with minimal within-individual variation over the breeding season (mean range = 4.6 ± 2.7%). In all individuals (n = 7), irregular iris pigmentation remained largely stable over the 2024 breeding season, indicating that once established, these changes seem to be maintained over time. This low within-individual variability strengthens the potential utility of iris pigmentation irregularities as a tool to document prior AIV infections, as we observed very little fading of irregular irises within a field season. Variability among individuals may still reflect differences in the magnitude of post-infection iris changes, potentially linked to immune recovery, timing of infection relative to sampling, or disease severity. Since all data were collected during the 2023 and 2024 seasons, well after the peak 2022 HPAIV outbreak, the overall stability observed suggests that iris pigmentation irregularities can persist for many months and may remain a useful marker of prior infection across breeding seasons. Nonetheless, we cannot rule out the possibility of continued or renewed exposure during the winter or early 2023. Supporting this, an antibody study in eggs from the same colony detected AIV-related antibodies in samples from 2023, suggesting that some females were either re-exposed during the pre-breeding period that year or retained antibodies from infections in 2022 (McLaughlin et al., 2025). While our findings indicate an association between iris pigmentation irregularities and AIV exposure, further studies are required to confirm a causal link and to evaluate how long these changes persist post-infection and whether they reflect exposure timing or severity.

### 4.3. Implications for Field Detection and Population Monitoring

While mortality has been the most visible consequence of H5 HPAIV clade 2.3.4.4b outbreaks in many seabird populations globally, the potential for sublethal effects is increasingly recognized as critical to understanding the full impact of infection and disease on wild populations. Sublethal consequences of AIV infections in birds have been reported in other systems, including reduced breeding propensity and movement (Gamble, 2023), delayed migration (Teitelbaum et al., 2023a; 2023b), lower body mass (Kuiken, 2013), and increased contaminant burdens (Teitelbaum et al., 2022). However, such effects are rarely documented in detail, particularly in free-ranging birds. In this context, our study provides empirical evidence linking a non-lethal phenotypic change to a previous viral exposure. Because iris pigmentation irregularities are relatively easy to observe in the field in gannets, they represent a low-cost, practical, and minimally invasive approach for detecting prior viral exposure, with potential for broad application in population monitoring. Given the visibility of these changes and their relatively stable appearance over the breeding season in our small sample, iris scoring could complement serological monitoring to assess past infection status. Our categorical analysis supports this potential: birds with moderate to high irregularity (>33% across both eyes) were more likely to be antibody positive than those with low irregularity (Figure 6). In the field, such a tool could be implemented through direct visual scoring during banding or nest checks, or via standardized photography. Processing images to quantify iris pigment irregularities is relatively fast, taking 5–10 minutes per iris with Photoshop, and could be further streamlined using free, open-source alternatives (e.g., ImageJ, https://imagej.net/ij/). However, environmental factors such as lighting conditions and viewing distance, as well as observer variability, may affect detection accuracy. Repeatability tests, inter-observer reliability, and standardized protocols will be essential for field application. Importantly, this method should be implemented in conjunction with traditional viral surveillance to provide a contextual understanding of local and temporal infection dynamics, which is necessary for interpreting the prevalence and meaning of observed pigmentation irregularities.

Iris pigment irregularities in our study were not thought to be associated with impaired behavior. Although we did not assess visual function directly, there was no evidence that these ocular changes impaired vision; birds with irregular iris pigmentation exhibited behaviour comparable to those with regular iris pigmentation from a distance before capture, during capture, and while being handled in the field (normal flight, foraging, and alert responses; Y.S., *personal communication*). Furthermore, we did not detect any relationship between anti-AIV antibody levels and body mass, and all individuals sampled appeared to be in good condition.

This suggests that variation in iris pigment irregularity is unlikely to impair foraging capacity through vision and therefore may not directly affect condition. While birds with irregular iris pigmentation were less often associated with active nests containing chicks, this pattern likely reflects our sampling design, as individuals were captured based on visible iris pigment irregularities regardless of nesting status, rather than a causal relationship. Although causality cannot be inferred, this pattern raises the possibility that pronounced iris pigmentation irregularities may coincide with reduced reproductive output. A complementary study by Lewis et al. (2025) provides important context for this interpretation. They found that gannets with black irises had similar breeding success to unaffected individuals at two United Kingdom colonies, suggesting that reproductive output among survivors is not substantially compromised. Together with our findings, this supports the potential for iris pigmentation irregularities to serve as a rapid, non-invasive indicator for post-outbreak population monitoring. Whereas Lewis et al. (2025) inferred prior exposure from iris colour alone, our study empirically validates that association through serological data, reinforcing the value of iris assessment as a scalable, field-based tool for tracking population recovery following disease-driven mortality events. Moreover, observations of the seven individuals monitored over the breeding season indicated generally successful reproduction: approximately 71% of nests (5 of 7) successfully produced chicks that survived to at least until our last observation in late July–early September, despite average irregularity in iris pigmentation all above 36%. The remaining nests were abandoned or empty at later visits, indicating that factors other than iris pigmentation may have influenced reproductive outcomes. Further investigation is warranted to assess potential fitness consequences and determine whether these reflect lingering physiological impacts or behavioral shifts post-infection.

### 4.4. Limitations and Future Directions

While our findings are consistent with a post-outbreak effect, we cannot conclusively determine whether iris pigmentation irregularities were driven specifically by H5 HPAIV, or another AIV strain, due to limitations in serotype resolution and sample size. Anti-NP antibodies provide evidence of general AIV exposure, while anti-H5 antibodies indicate exposure to an H5 subtype but not necessarily H5 HPAIV. However, given the colony’s negative status prior to 2022 (Lane et al., 2023), the confirmed detection of H5 HPAIV during large-scale mortality events, and the lack of evidence for concurrent circulation of low-path H5 or other AIV strains, we assume that most antibody-positive individuals were exposed to H5 HPAIV. We acknowledge the limitations of this assumption and recommend further virological and serological surveillance to refine this interpretation. Further, we observed a weak but statistically significant positive correlation between iris irregularities and anti-H5 antibody levels. However, irregularity was not a significant predictor of anti-H5 antibody status (i.e., positive/negative), indicating that the extent of iris pigmentation irregularity does not consistently reflect current H5 antibody detectability. This discrepancy between H5 and NP results may reflect different waning rates of antibody types, as reported in other wild bird studies (Giacinti et al., 2024b; Wight, 2024), with NP antibodies typically persisting longer than subtype-specific H5 antibodies. This underscores the need for more precise temporal sampling. Additionally, Nobuto filter papers, particularly when stored at room temperature, are not considered the gold-standard method for antibody detection (Giacinti et al., 2025), and some discrepancies may reflect reduced assay sensitivity related to sample storage or elution efficiency.

Iris pigmentation changes may not be specific to AIV. Similar alterations could, in theory, arise from other infections or inflammatory processes that induce local immune responses or melanin deposition in the iris. Although we found no evidence of concurrent disease outbreaks at the colony, we cannot fully exclude alternative causes of inflammation or pigment change. Further, our analyses did not explicitly incorporate covariates such as sex, age, or body condition beyond body mass. These may influence immune response, disease recovery, or pigmentation patterns and should be considered in future models to clarify potential confounding effects (Wight, 2024).

While disease indicators with visible external manifestations have been documented in the past, such as bill deformities linked to avian keratin disorder (Van Hemert et al., 2012; 2013; Zylberberg et al., 2018) and ocular swelling from *Mycoplasma gallisepticum* (a prokaryote) infections (Thomason et al., 2017) in birds, these tend to be either transient or correlated with severe declines in fitness or survival. In contrast, few studies have identified subtle, non-debilitating, and quantifiable morphological traits, like iris pigmentation irregularities, that could reliably persist across time and reflect prior infection in wild populations. The lack of association between iris pigmentation irregularities and body mass, a proxy for apparent condition, suggests that these changes do not reflect ongoing physiological dysfunction. Instead, they may represent a sublethal, residual outcome of prior infection. This approach may be particularly valuable in pale-iris species, where pigmentation anomalies are more easily detected, and in contexts where repeated handling or blood sampling is impractical. However, we emphasize that the irregularity thresholds used here should be viewed as provisional. Future studies using larger sample sizes, cross-seasonal data, and populations with varying levels of exposure to AIV are needed to evaluate the diagnostic sensitivity, specificity, and temporal stability of iris changes as a biomarker.

The potential for iris pigmentation irregularity to change over time, whether via recovery or repeated exposures, presents both a strength and a challenge. While it may function as a dynamic biomarker, repeated infections could complicate interpretation. Thus, despite the promise of this approach, we caution that iris pigmentation may lose diagnostic value with repeated exposure. It remains unclear how multiple infections would influence iris changes, whether through progressive pigmentation, stabilization, or reversal. Also, once a bird develops iris irregularities, detecting subsequent exposures may be more difficult. We recommend that future field efforts include systematic annual iris photography or scoring of both eyes to build robust individual-level infection histories.

Overall, visual markers, like changes in iris pigmentation, are particularly valuable because they allow for non-invasive (once established with set thresholds), long-term monitoring of disease dynamics. These traits could be incorporated into field-based surveillance, offering a relatively low-cost and minimal-impact method for assessing past exposure at the population level. Together, our findings suggest that iris irregularities may represent a novel biomarker of prior AIV infection in gannets, expanding the toolkit available for wildlife disease ecology.

## CRediT authorship contribution statement

Christina Petalas: Methodology, Funding acquisition, Investigation, Formal analysis, Visualization, Writing – original draft, Writing – review & editing.

Yannick Seyer: Conceptualization, Methodology, Funding acquisition, Data acquisition, Investigation, Writing – review & editing

Jean-François Rail: Conceptualization, Methodology, Funding acquisition, Data acquisition, Investigation, Writing – review & editing

Jolene Giacinti: Methodology, Funding acquisition, Investigation, Writing – review & editing

Jennifer Provencher: Methodology, Funding acquisition, Investigation, Writing – review & editing

Kyle Elliott: Funding acquisition, Writing – review & editing

Raphaël Lavoie: Methodology, Funding acquisition, Investigation, Writing – review & editing Stephanie Avery-Gomm: Funding acquisition, Writing – review & editing

## Data availability

The datasets generated and analyzed during the current study will be made publicly available on Open Government (https://open.canada.ca/en) at the time of publication. The data will be fully curated and documented to allow reproduction of all analyses presented in this paper.

## Acknowledgments

We would like to thank the Parc national de l’Île-Bonaventure-et-du-Rocher-Percé for its support in the field, and the field assistants who aided in data collection: Hilde Marie Johansen, Kevin Westeel, Anaïs Kerric, Léa Desjardins, Roxanne Turgeon, and Catherine Čapkun-Huot.

## References

Alexandrou, O., Catsadorakis, G. & U. Höfle. 2025. Ecoepidemiology of avian influenza and other diseases in Dalmatian pelicans – Summary Brief. Society for the Protection of Prespa, Greece/SaBio Health and Biotechnology Department, Institute for Game and Wildlife Research (UCLM-CSIC-JCCM), Spain. 36 pp. 10.5281/zenodo.15576290.

Avery Gomm, S., Barychka, T., English, M., Ronconi, R. A., Wilhelm, S. I., Rail, J. F., … & Wight, J. (2024). Wild bird mass mortalities in eastern Canada associated with the Highly Pathogenic Avian Influenza A (H5N1) virus, 2022. Ecosphere, 15(9), e4980.

Banyard, A. C., Lean, F. Z., Robinson, C., Howie, F., Tyler, G., Nisbet, C., … & Reid, S. M. (2022). Detection of highly pathogenic avian influenza virus H5N1 clade 2.3. 4.4 b in great skuas: a species of conservation concern in Great Britain. Viruses, 14(2), 212.

Blagodatski, A., Trutneva, K., Glazova, O., Mityaeva, O., Shevkova, L., Kegeles, E., … & Volchkov, P. (2021). Avian influenza in wild birds and poultry: dissemination pathways, monitoring methods, and virus ecology. Pathogens, 10(5), 630.

Brown, J. D., Stallknecht, D. E., Berghaus, R. D., Luttrell, M. P., Velek, K., Kistler, W., Costa, T., Yabsley, M. J., & Swayne, D. (2009). Evaluation of a Commercial Blocking Enzyme-Linked Immunosorbent Assay to Detect Avian Influenza Virus Antibodies in Multiple Experimentally Infected Avian Species. Clinical and Vaccine Immunology, 16(6), 824–829. 10.1128/CVI.00084-09

Camphuysen, C. J., Gear, S. C., & Furness, R. W. (2022). Avian influenza leads to mass mortality of adult Great Skuas in Foula in summer 2022. Scott. Birds, 42, 312–323.

Careen, N. G., Collins, S. M., D’entremont, K. J., Wight, J., Rahman, I., Hargan, K. E., … & Montevecchi, W. A. (2024). Highly pathogenic avian influenza virus resulted in unprecedented reproductive failure and movement behaviour by northern gannets. Marine Ornithology, 52(1).

Chatziprodromidou, I. P., Arvanitidou, M., Guitian, J., Apostolou, T., Vantarakis, G., & Vantarakis, A. (2018). Global avian influenza outbreaks 2010–2016: A systematic review of their distribution, avian species and virus subtype. Systematic reviews, 7, 1–12.

CMS FAO Co-convened Scientific Task Force on Avian Influenza and Wild Birds (2022). Scientific Task Force on Avian Influenza and Wild Birds statement. H5N1 Highly Pathogenic Avian Influenza in poultry and wild birds: Winter of 2021/2022 with focus on mass mortality of wild birds in UK and Israel. Available at: https://www.cms.int/sites/default/files/uploads/avian_influenza_0.pdf

Corbett, E. C., Brumfield, R. T., & Faircloth, B. C. (2024). The mechanistic, genetic and evolutionary causes of bird eye colour variation. Ibis, 166(2), 560–589.

Dewar, M., Wille, M., Gamble, A., Vanstreels, R. E., Bouliner, T., Smith, A., … & Hart, T. (2023). The risk of highly pathogenic avian influenza in the Southern Ocean: a practical guide for operators and scientists interacting with wildlife. Antarctic Science, 35(6), 407–414.

Duriez, O., Sassi, Y., Le Gall-Ladevèze, C., Giraud, L., Straughan, R., Dauverné, L., … & Le Loc’h, G. (2023). Highly pathogenic avian influenza affects vultures’ movements and breeding output. Current Biology, 33(17), 3766–3774.

Gamble, A. (2023). Disease ecology: When a GPS logger tells you more than a blood sample. Current Biology, 33(17), R907–R909.

Giacinti, J. A., Jarvis-Cross, M., Lewis, H., Provencher, J. F., Berhane, Y., Kuchinski, K., … & Sharp, C. M. (2024b). Transmission dynamics of highly pathogenic avian influenza virus at the wildlife-poultry-environmental interface: A case study. One Health, 19, 100932.

Giacinti, J. A., Signore, A. V., Jones, M. E., Bourque, L., Lair, S., Jardine, C., … & Soos, C. (2024a). Avian influenza viruses in wild birds in Canada following incursions of highly pathogenic H5N1 virus from Eurasia in 2021–2022. Mbio, 15(8), e03203–23.

Giacinti, J.A., Rahman, I., Wight, J., Lewis, H., Taylor, L.U., Provencher, J.F., Ronconi, R., Berhane, Y., Xu, W., Zhmendak, D., Sarma, S.N., Sharp, C.M., Cunningham, J.T., Hedd, A., Bosch, J., Robertson, G.J., Hargan, K.E., Lang, A.S. (2025). Comparison of whole blood on filter strips with serum for avian influenza virus antibody detection in wild birds. Conservation Physiology, 13(1), coaf033.

Grémillet, D., Péron, C., Lescroël, A., Fort, J., Patrick, S. C., Besnard, A., & Provost, P. (2020). No way home: collapse in northern gannet survival rates point to critical marine ecosystem perturbation. Marine Biology, 167(12), 189.

Guillemette, M., Grégoire, F., Bouillet, D., Rail, J. F., Bolduc, F., Caron, A., & Pelletier, D. (2018). Breeding failure of seabirds in relation to fish depletion: Is there one universal threshold of food abundance?. Marine Ecology Progress Series, 587, 235–245.

Hochman, O., Xu, W., Yang, M., Yang, C., Ambagala, A., Rogiewicz, A., Wang, J. J., & Berhane, Y. (2023). Development and Validation of Competitive ELISA for Detection of H5 Hemagglutinin Antibodies. Poultry, 2(3), 349–362. 10.3390/poultry2030026

Hoye, B. J., Munster, V. J., Nishiura, H., Klaassen, M., & Fouchier, R. A. (2010). Surveillance of wild birds for avian influenza virus. Emerging infectious diseases, 16(12), 1827.

James, J., Seekings, A. H., Skinner, P., Purchase, K., Mahmood, S., Brown, I. H., … & Reid, S. M. (2022). Rapid and sensitive detection of high pathogenicity Eurasian clade 2.3. 4.4 b avian influenza viruses in wild birds and poultry. Journal of virological methods, 301, 114454.

Jessopp, M., Giralt Paradell, O., Goh, T., Popov, D., & Rogan, E. (2023). Estimated mortality of the highly pathogenic avian influenza pandemic on northern gannets (Morus bassanus) in southwest Ireland.

Knief, U., Bregnballe, T., Alfarwi, I., Ballmann, M. Z., Brenninkmeijer, A., Bzoma, S., … & Courtens, W. (2024). Highly pathogenic avian influenza causes mass mortality in Sandwich Tern Thalasseus sandvicensis breeding colonies across north-western Europe. Bird conservation international, 34, e6.

Kuiken, T. (2013). Is low pathogenic avian influenza virus virulent for wild waterbirds?. Proceedings of the Royal Society B: Biological Sciences, 280(1763), 20130990.

Lane, J. V., Jeglinski, J. W., Avery Gomm, S., Ballstaedt, E., Banyard, A. C., Barychka, T., … & Votier, S. C. (2023). High pathogenicity avian influenza (H5N1) in Northern Gannets (Morus bassanus): Global spread, clinical signs and demographic consequences. Ibis, 166(2), 633–650.

Lang, A. S., Lebarbenchon, C., Ramey, A. M., Robertson, G. J., Waldenström, J., & Wille, M. (2016). Assessing the role of seabirds in the ecology of influenza A viruses. Avian diseases, 60(1s), 378–386.

Lewis, S., Burton, E., Butcher, J., Cleasby, I., King, A., Marriott, E., … & Lane, J. V. (2025). Effect of a previous high pathogenicity avian influenza (HPAIV) infection on the breeding success of Northern Gannets (Morus bassanus). Ibis.

Lycett, S. J., Duchatel, F., & Digard, P. (2019). A brief history of bird flu. Philosophical Transactions of the Royal Society B, 374(1775), 20180257.

McLaughlin, A., Giacinti, J., Sarma, S. N., Brown, M. G., Ronconi, R. A., Lavoie, R. A., … & Provencher, J. F. (2025). Examining avian influenza virus exposure in seabirds of the northwest Atlantic in 2022 and 2023 via antibodies in eggs. Conservation Physiology, 13(1), coaf010.

McPhail, G. M., Collins, S. M., Burt, T. V., Careen, N. G., Doiron, P. B., Avery-Gomm, S., … & Montevecchi, W. A. (2024). Geographic, ecological, and temporal patterns of seabird mortality during the 2022 HPAI H5N1 outbreak on the island of Newfoundland. Canadian Journal of Zoology, 103, 1–12.

Montevecchi, W. A., Regular, P. M., Rail, J. F., Power, K., Mooney, C., D’entremont, K. J., … & Wilhelm, S. I. (2021). Ocean heat wave induces breeding failure at the southern breeding limit of the Northern Gannet Morus bassanus. Marine Ornithology, 49, 71–78.

Mowbray, T. B. (2020). Northern Gannet (Morus bassanus), version 1.0. Birds of the World (SM Billerman, Editor). Cornell Lab of Ornithology, Ithaca, NY, USA. doi, 10.

Nagy, A., Černíková, L., Kunteová, K., Dirbáková, Z., Thomas, S. S., Slomka, M. J., … & Brown, I. H. (2021). A universal RT-qPCR assay for “One Health” detection of influenza A viruses. PloS one, 16(1), e0244669.

Nalepa, R., Provencher, J., Giacinti, J. A., Wilcox, A., Sharp, C. M., Ronconi, R. A., … & Avery-Gomm, S. (2024). An expert opinion process to prioritize One Health information needs during a zoonotic disease outbreak. FACETS.

Olson, V. A., & Owens, I. P. F. (2005). Interspecific variation in the use of carotenoid based coloration in birds: diet, life history and phylogeny. Journal of Evolutionary Biology, 18(6), 1534–1546.

Pohlmann, A., Stejskal, O., King, J., Bouwhuis, S., Packmor, F., Ballstaedt, E., … & Harder, T. (2023). Mass mortality among colony-breeding seabirds in the German Wadden Sea in 2022 due to distinct genotypes of HPAIV H5N1 clade 2.3. 4.4 b. Journal of General Virology, 104(4), 001834.

Poulin, J. M. (1968). Reproduction du Fou de Bassan (Sula bassana), Ile Bonaventure (Québec). Master’s Thesis, Univ. Laval, Québec.

R Core Team. (2025). R: A language and environment for statistical computing. R Foundation for Statistical Computing, Vienna, Austria. https://www.R-project.org/

Rail, J. F. (2021). Eighteenth census of seabirds breeding in the sanctuaries of the North Shore of the Gulf of St. Lawrence, 2015. The Canadian Field-Naturalist, 135(3), 221–233.

Rijks, J. M., Leopold, M. F., Kühn, S., Schenk, F., Brenninkmeijer, A., Lilipaly, S. J., … & Beerens, N. (2022). Mass mortality caused by highly pathogenic influenza A (H5N1) virus in sandwich terns, the Netherlands, 2022. Emerging Infectious Diseases, 28(12), 2538.

Seyer, Y. and M. Guillemette. (2023). Situation of the Northern Gannet colony of Bonaventure Island (QC) following the outbreak of highly pathogenic avian influenza (HPAI) virus in the spring of 2022 in eastern Canada. Report. 38 + xii pages. Université du Québec à Rimouski. Rimouski, Québec, Canada.

Siebert, U., Schwemmer, P., Guse, N., Harder, T., Garthe, S., Prenger-Berninghoff, E., & Wohlsein, P. (2012). Health status of seabirds and coastal birds found at the German North Sea coast. Acta Veterinaria Scandinavica, 54(1), 43.

Stallknecht, D. E., & Brown, J. D. (2008). Ecology of avian influenza in wild birds. Avian influenza, 43–58.

Teitelbaum, C. S., Ackerman, J. T., Hill, M. A., Satter, J. M., Casazza, M. L., De La Cruz, S. E., … & Prosser, D. J. (2022). Avian influenza antibody prevalence increases with mercury contamination in wild waterfowl. Proceedings of the Royal Society B, 289(1982), 20221312.

Teitelbaum, C. S., Casazza, M. L., McDuie, F., De La Cruz, S. E., Overton, C. T., Hall, L. A., … & Prosser, D. J. (2023a). Waterfowl recently infected with low pathogenic avian influenza exhibit reduced local movement and delayed migration. Ecosphere, 14(2), e4432.

Teitelbaum, C. S., Masto, N. M., Sullivan, J. D., Keever, A. C., Poulson, R. L., Carter, D. L., … & Prosser, D. J. (2023b). North American wintering mallards infected with highly pathogenic avian influenza show few signs of altered local or migratory movements. Scientific reports, 13(1), 14473.

Thomason, C. A., Leon, A., Kirkpatrick, L. T., Belden, L. K., & Hawley, D. M. (2017). Eye of the Finch: characterization of the ocular microbiome of house finches in relation to mycoplasmal conjunctivitis. Environmental microbiology, 19(4), 1439–1449.

Toomey, M. B., Butler, M. W., & McGraw, K. J. (2010). Immune-system activation depletes retinal carotenoids in house finches (Carpodacus mexicanus). Journal of Experimental Biology, 213(10), 1709–1716.

Tremlett, C. J., Morley, N., & Wilson, L. J. (2024). UK seabird colony counts in 2023 following the 2021–22 outbreak of Highly Pathogenic Avian Influenza. RSPB Research Report 76.

Van Hemert, C., Armién, A. G., Blake, J. E., Handel, C. M., & O’Hara, T. M. (2013). Macroscopic, histologic, and ultrastructural lesions associated with avian keratin disorder in black-capped chickadees (Poecile atricapillus). Veterinary pathology, 50(3), 500–513.

Van Hemert, C., Handel, C. M., & O’Hara, T. M. (2012). Evidence of accelerated beak growth associated with avian keratin disorder in black-capped chickadees (Poecile atricapillus). Journal of Wildlife Diseases, 48(3), 686–694.

Wight, J. (2024). Wild bird mass mortalities in eastern Canada associated with the Highly Pathogenic Avian Influenza A (H5N1) virus, 2022. Ecosphere, 15(9), e4980.

Wille, M., & Barr, I. G. (2022). Resurgence of avian influenza virus. Science, 376(6592), 459–460.

Williams, S. M., Huwiler, M., & Mehlhorn, H. (2001). Ocular and encephalic toxoplasmosis in canaries (Serinus canaria). Avian Diseases, 45(4), 1134–1137. 10.1637/0005-2086(2001)045[1134:OETIC]2.0.CO;2

Yamamoto, Y., Nakamura, K., Yamada, M., & Mase, M. (2016). Corneal opacity in domestic ducks experimentally infected with H5N1 highly pathogenic avian influenza virus. Veterinary Pathology, 53(1), 65–76.

Zylberberg, M., Van Hemert, C., Handel, C. M., & DeRisi, J. L. (2018). Avian keratin disorder of Alaska black-capped chickadees is associated with Poecivirus infection. Virology Journal, 15, 1–9.

