## Supplemental Materials for "IRIS PIGMENTATION IRREGULARITIES FOLLOWING AN AVIAN INFLUENZA OUTBREAK: IMPLICATIONS FOR DISEASE SURVEILLANCE AND POPULATION MONITORING IN A COLONIAL SEABIRD"

^e^Wildlife Research Division, Wildlife and Landscape Science Directorate, Science and Technology Branch, Environment and Climate Change Canada, Ottawa, Ontario, Canada

^f^Ecotoxicology and Wildlife Health Division, Wildlife and Landscape Science Directorate, Science and Technology Branch, Environment and Climate Change Canada, Québec, Québec Canada

**This supporting file includes: 6 pages, 4 figures**

**Table of Contents**

|  | Description | Page |
| --- | --- | --- |
| Figure S1 | Correlation between anti-NP S/N values and iris pigmentation irregularity (%) for the iris with greater irregularity. Points are colored by anti-NP result interpretations (“Positive” or “Negative”). The dashed light green line shows the average pre-outbreak (pre-2021) irregularity, with the shaded band representing ±1 standard deviation. Note that lower S/N ratios correspond to stronger anti-NP antibody detection. An optimized threshold of S/N < 0.77 was used to classify samples as positive for anti-NP antibodies, based on threshold optimization for Nobuto strip testing at NWRC (Giacinti et al., 2025). | S3 |
| Figure S2 | Correlation between anti-H5 PI% antibody detection and iris pigmentation irregularity (%) for the iris with greater irregularity. Points are colored by anti-H5 result interpretations (“Positive” or “Negative”). The dashed light green line shows the average pre-outbreak (pre-2021) irregularity, with the shaded band representing ±1 standard deviation. Note that higher PI values correspond to stronger anti-H5 antibody detection. A PI value ≥ 20.37% was considered positive, reflecting the optimized cutoff for Nobuto strips (Giacinti et al., 2025). | S4 |
| Figure S3 | Logistic regression models predicting the probability of anti-NP S/N antibody positivity based on the iris with greater irregularity. Points are colored by test result interpretation (“Positive” or “Negative”) for each sample. The dashed light green line is average pre-outbreak (pre-2021) calculated irregularity, with the shaded band representing ±1 standard deviation. The solid blue line indicates the predicted probability of seropositivity based on the fitted regression model. | S5 |
| Figure S4 | Example of a Northern Gannet exhibiting minimal change in iris pigmentation irregularity throughout the 2024 breeding season on Bonaventure Island. | S6 |

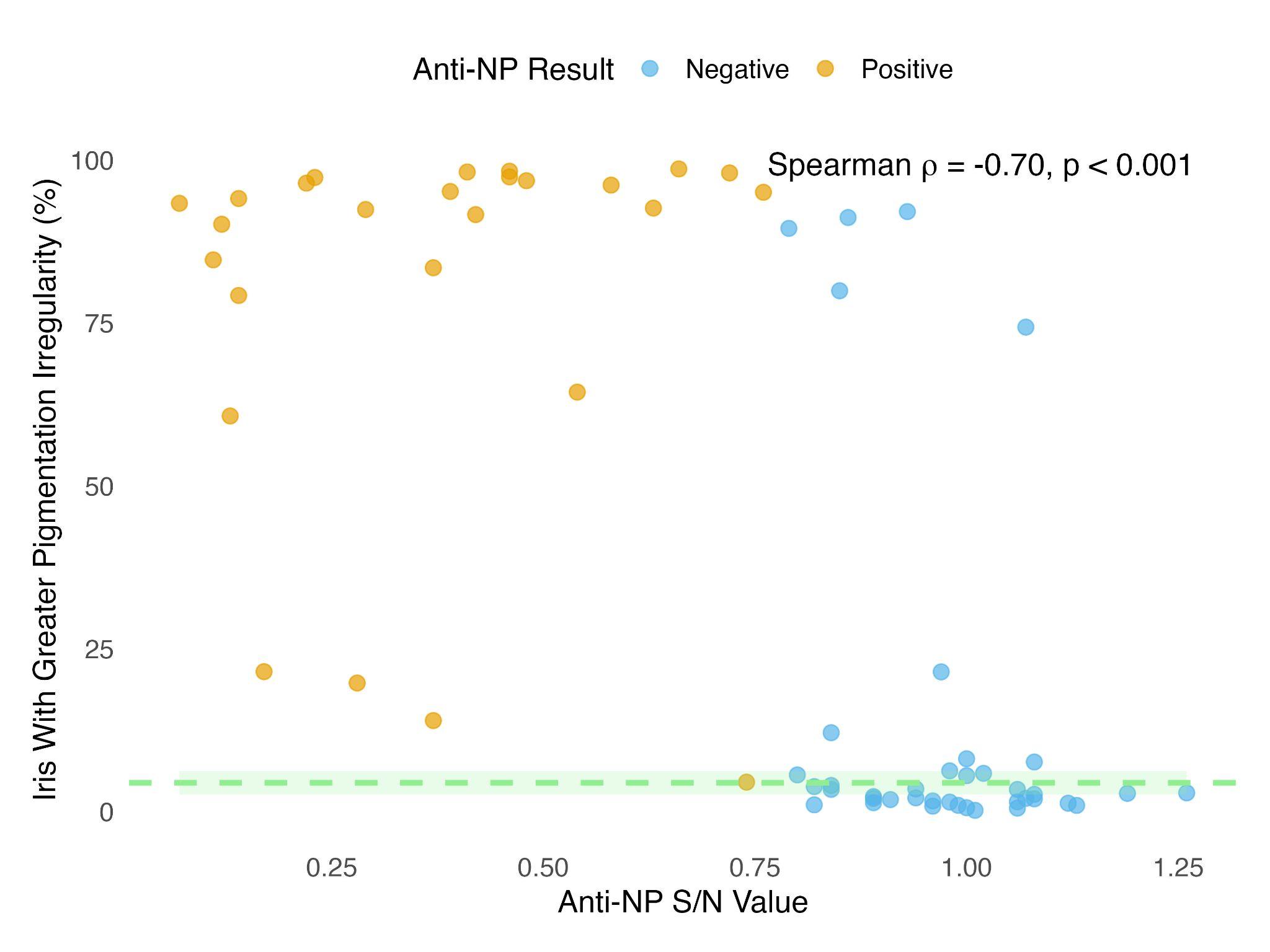

**Figure S1.** Correlation between anti-NP S/N values and iris pigmentation irregularity (%) for the iris with greater irregularity. Points are colored by anti-NP result interpretations (“Positive” or “Negative”). The dashed light green line shows the average pre-outbreak (pre-2021) irregularity, with the shaded band representing ±1 standard deviation. Note that lower S/N ratios correspond to stronger anti-NP antibody detection. An optimized threshold of S/N < 0.77 was used to classify samples as positive for anti-NP antibodies, based on threshold optimization for Nobuto strip testing at NWRC (Giacinti et al., 2025).

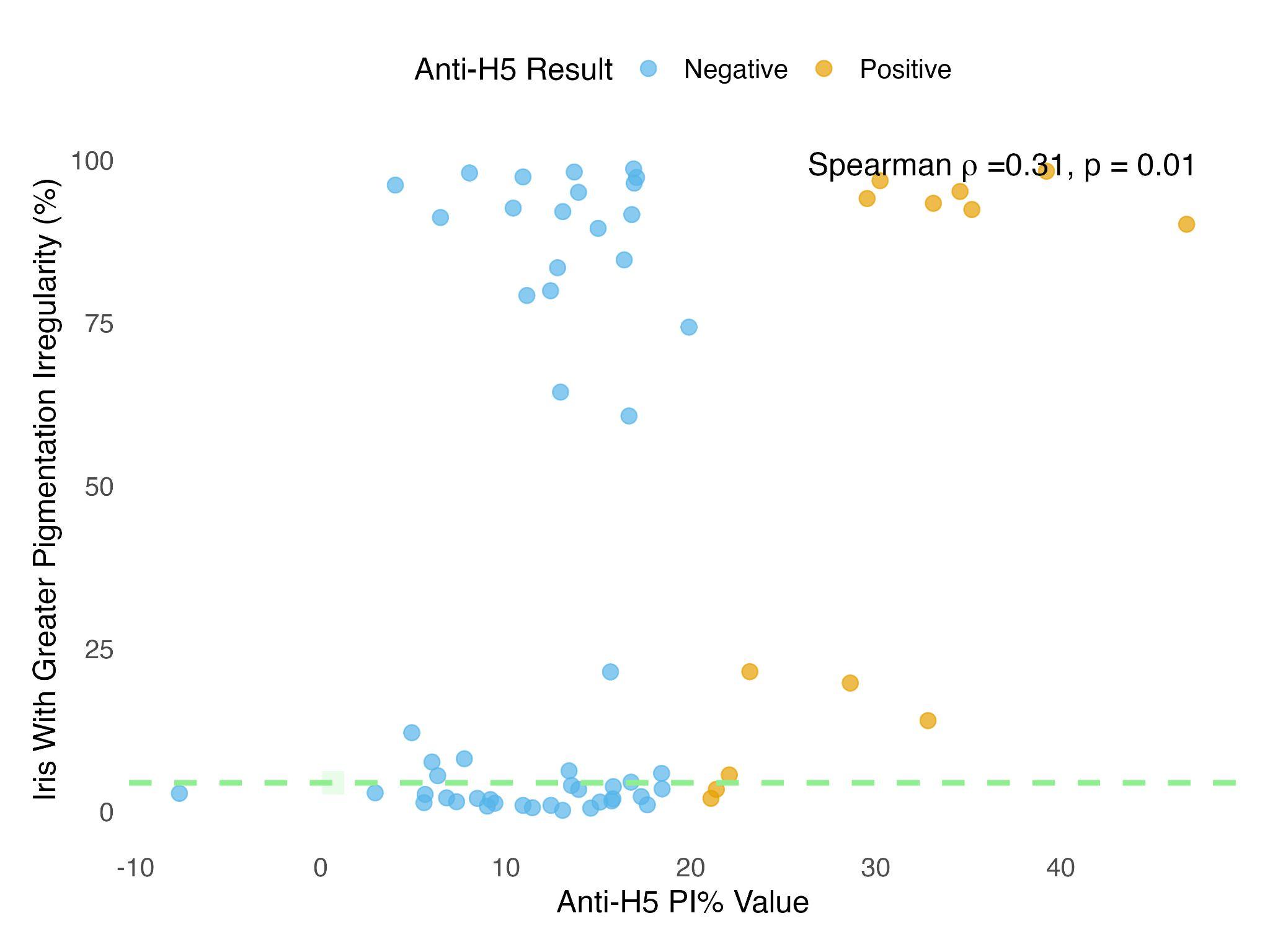

**Figure S2.** Correlation between anti-H5 PI% antibody detection and iris pigmentation irregularity (%) for the iris with greater irregularity. Points are colored by anti-H5 result interpretations (“Positive” or “Negative”). The dashed light green line shows the average pre-outbreak (pre-2021) irregularity, with the shaded band representing ±1 standard deviation. Note that higher PI values correspond to stronger anti-H5 antibody detection. A PI value ≥ 20.37% was considered positive, reflecting the optimized cutoff for Nobuto strips (Giacinti et al., 2025).

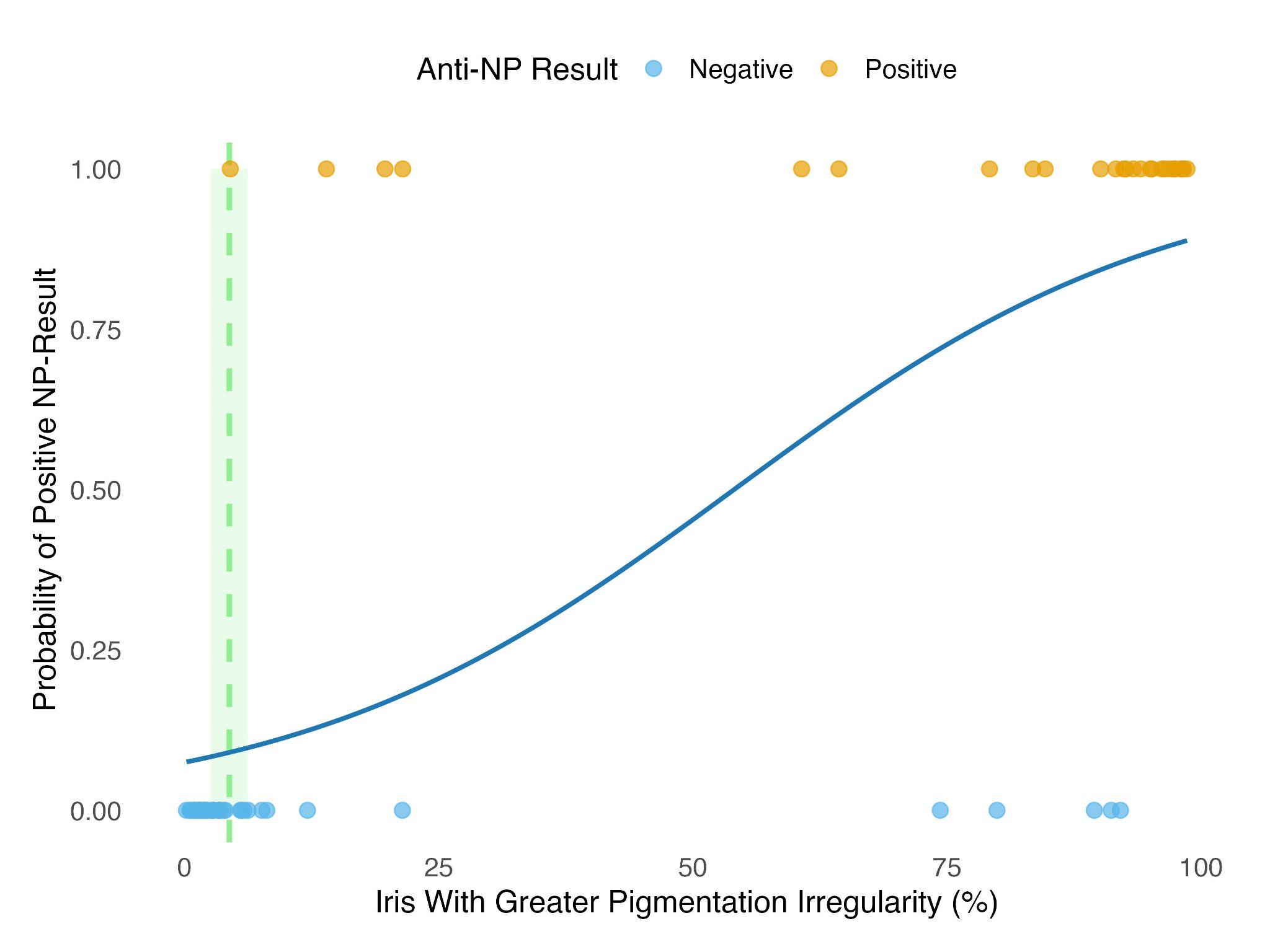

**Figure S3.** Logistic regression models predicting the probability of anti-NP S/N antibody positivity based on the iris with greater irregularity. Points are colored by test result interpretation (“Positive” or “Negative”) for each sample. The dashed light green line is average pre-outbreak (pre-2021) calculated irregularity, with the shaded band representing ±1 standard deviation. The solid blue line indicates the predicted probability of seropositivity based on the fitted regression model.

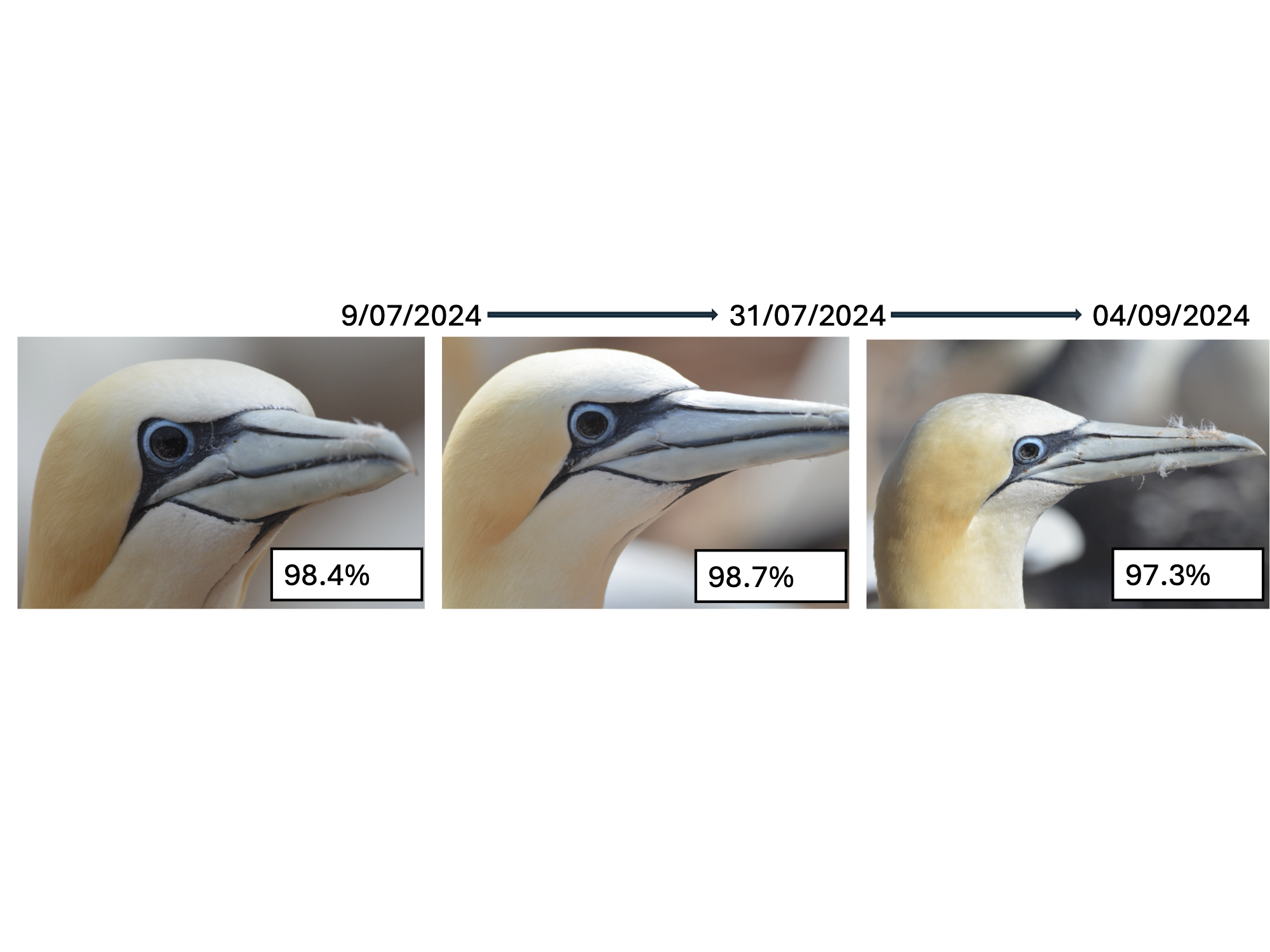

**Figure S4.** Example of a Northern Gannet exhibiting minimal change in iris pigmentation irregularity throughout the 2024 breeding season on Bonaventure Island. The percentage represents the calculated iris pigmentation irregularity for the eye being pictured.
